# Epigenetic reprogramming of imprinting at meiosis

**DOI:** 10.1101/2023.05.17.541143

**Authors:** Sean A. Montgomery, Frédéric Berger

## Abstract

In mammals, genomic imprinting results from different sets of epigenetic marks that distinguish the parental origins of loci in the progeny. Epigenetic reprogramming of genomic imprinting is necessary to establish a totipotent cell state. The consecutive erasure of parental epigenetic marks and the deposition of new marks occurs alongside major life stage transitions including gametogenesis and fertilization. However, despite occurring concomitantly with gametogenesis, the role of meiosis in epigenetic reprogramming has received little attention. To address this question, we use the model bryophyte *Marchantia polymorpha*. Following the haploid reproductive phase of this land plant, the expression of the paternal genome is silenced by the histone modification H3K27me3 in the short-lived diploid embryo. We show that imprinting is erased during meiosis, which occurs separately from gametogenesis and fertilization in Marchantia. The epigenetic reprogramming initiated during meiosis is completed in the meiotic spores where the chromatin landscape of the next haploid generation is established *de novo*. Hence, our findings illustrate a potential role for meiosis in epigenetic reprogramming that may be generalized to other sexually reproducing species.

## Introduction

Epigenetic reprogramming erases the parental epigenomes inherited from the gametes and establishes the chromatin landscape of the progenitor of a new generation. In animals, a key role for reprogramming is the establishment of totipotency of the single-celled zygote via the germ cells to allow it to produce all cell types within an organism (Farhadova et al., 2019; Gurdon, 2017; Hemberger et al., 2009; Irie et al., 2018). Failures in reprogramming result in disease (Tucci et al., 2019). In placental mammals, one major barrier to reprogramming is genomic imprinting, the asymmetric distribution of chromatin modifications between parental alleles resulting in biased gene expression (Barlow and Bartolomei, 2014; Ferguson-Smith, 2011; Pires and Grossniklaus, 2014; Tucci et al., 2019). While DNA methylation constitutes the epigenetic mark for many imprinted loci, repressive histone post-translational modifications (PTMs) and non-coding RNAs are also involved in the control of imprinting (Bartolomei and Ferguson-Smith, 2011; Inoue, 2023; Llères et al., 2021; SanMiguel and Bartolomei, 2018). Erasing existing genomic imprints and establishing the proper epigenetic landscape is achieved naturally around the time of gametogenesis, meiosis, and fertilization (Kelsey and Feil, 2013; Luo et al., 2018; Nashun et al., 2015). However, the overlapping timing of germ cell differentiation, meiosis, and fertilization hinders parsing out the relative impact of each process on reprogramming (Kota and Feil, 2010).

Genomic imprinting has been described outside of placental mammals, permitting the impact of gamete differentiation, fertilization, or meiosis on reprogramming to be studied independent of one another. Extensive studies in flowering plants have revealed imprinted loci to be confined to the endosperm, an extra-embryonic tissue analogous to the placenta (Batista and Köhler, 2020; Gehring, 2013). However, the endosperm does not contribute to the next generation (Gehring and Satyaki, 2017), thus reprogramming of imprinting is not required and unlikely in flowering plants. Other forms of imprinting affect not only isolated genes, but entire chromosomes, for example paternal X chromosome inactivation in mammals (Żylicz and Heard, 2020) and the heterochromatinization of paternal chromosomes in insects (Hodson and Ross, 2021) and lampreys (Timoshevskaya et al., 2023). In the model bryophyte Marchantia, Paternal Chromosome Repression (PCR) is a unique form of genomic imprinting in which the entire paternal genome is silenced in the embryo by a histone PTM, trimethylation of lysine 27 on histone H3 (H3K27me3), resulting in gene expression primarily from maternal alleles (Montgomery et al., 2022). Like all land plants, the life cycle of Marchantia alternates between haploid and diploid generations but, unlike flowering plants, vegetative development is haploid and initiated by germination of a spore (Fig. 1A) (Shimamura, 2015). Gametogenesis and fertilization generate the diploid embryo (also called sporophyte). After the establishment of genomic imprinting in diploid embryos, most of the embryonic cells undergo meiosis. Meiosis generates spores, which are totipotent and develop into the haploid adult plant (Fig. 1A). Therefore, unlike in mammals and flowering plants, meiosis is clearly separated from gametogenesis and fertilization. We reasoned that Marchantia would therefore be an ideal model to study the impact of meiosis on the reprogramming of genomic imprinting.

**Figure 1.**
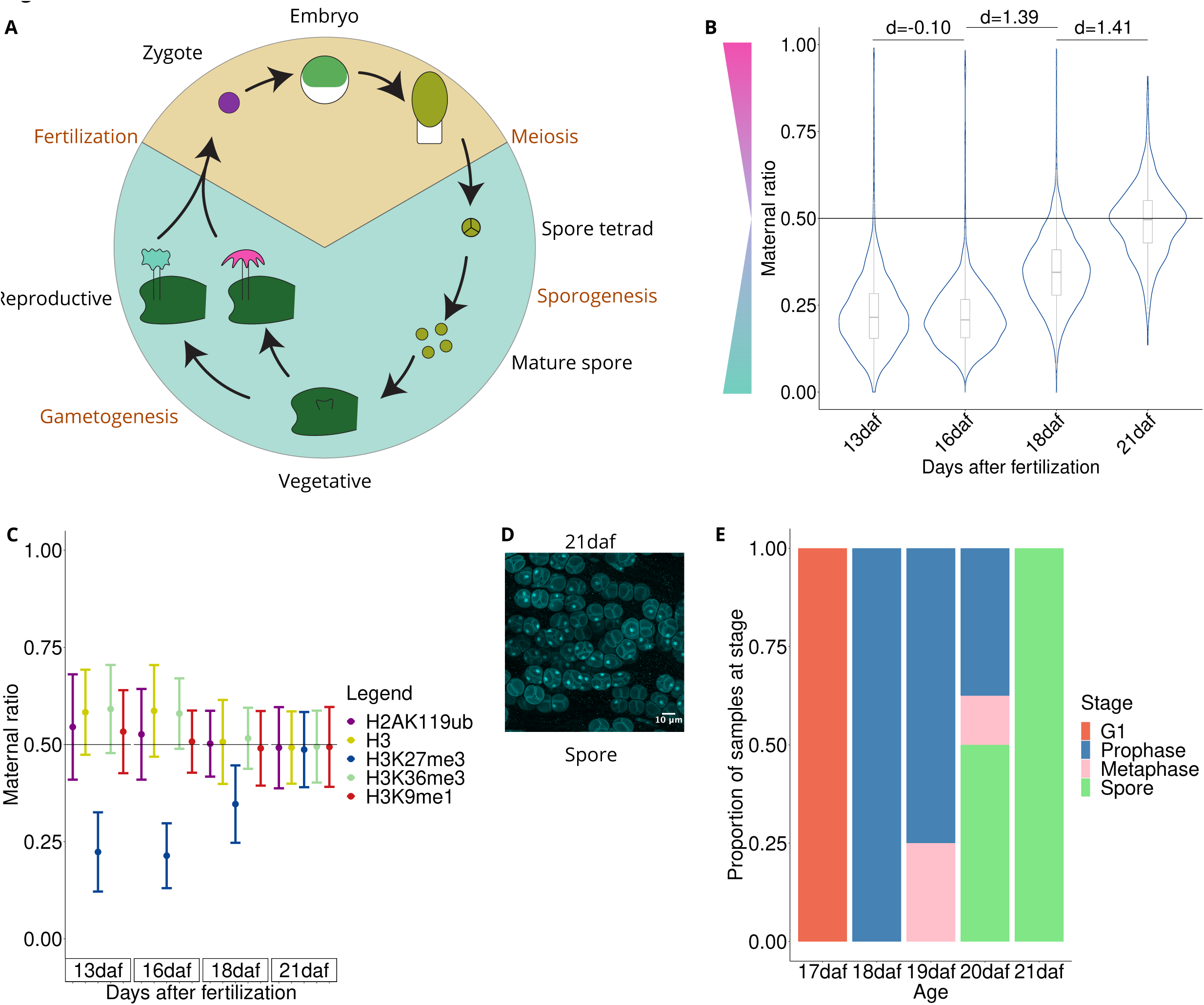
Loss of imprinting after meiosis. (A) Diagram of the Marchantia life cycle. Cyan and brown backgrounds indicate haploid and diploid life cycle phases, respectively. Black text denotes developmental stages and orange text denotes developmental transitions. (B) Violin plot of maternal ratio of H3K27me3 per gene. daf = days after fertilization. Cohen’s *d* effect size values are indicated for pairwise comparisons of consecutive stages, where |*d*| < 0.2 is a negligible effect and 0.8 < |*d*| is a large effect, as previously reported (Cohen, 1992). (C) Maternal ratio of all chromatin marks profiled. Dots indicate mean maternal ratio per gene per mark and lines indicate standard deviation. (D) Representative DAPI staining of 21 daf sample. (E) Quantification of developmental stages as pre-meiosis (G1), meiotic (Prophase and Metaphase), and post-meiotic spore tetrads (Spore) from individual sporophytes (N=8 for each stage).

Here, we investigated the dynamics of histone PTMs through embryo development and meiosis, including the imprinting mark H3K27me3, which is responsible for PCR, and other histone PTMs that do not participate in imprinting yet regulate transcription. We established the erasure of genome-wide imprints of the embryo by completion of meiosis, followed by the establishment of the chromatin landscape of the haploid vegetative plant in meiotic spores. Hence, meiosis is associated with two steps of reprogramming of the epigenetic landscape of the diploid embryo that prepares chromatin organization in spores for the initiation of the new life cycle.

## Results

### PCR is not dependent on accessions investigated and does not involve DNA methylation

In our previous study, we analyzed crosses between female Cam-2 and male Tak-1 as well as between Tak-2 and Tak-1 (Montgomery et al., 2022). To investigate the robustness of PCR across Marchantia accessions, we performed additional crosses and generated transcriptomes from embryos. To use an accession distinct from the Tak-1 male in crosses, we sequenced the genome of the male Cam-1 accession, performed crosses between Cam-2 and Cam-1, and generated transcriptomes from embryos. After assigning reads to maternal or paternal genomes on the basis of single-nucleotide polymorphisms (SNPs) between accessions, we calculated for each gene a maternal ratio (*p_m_*) of reads which ranges from 0 to 1, where 0 represents detection of only paternal reads and 1 represents detection of only maternal reads. Transcription was generally maternally biased, with 95% of genes classified as having maternally biased or fully maternal transcription (Fig. S1A-B). We conclude that PCR does not depend on Tak-1 males but is likely general across Marchantia natural accessions.

Cytosine DNA methylation is often associated with imprinting (Ferguson-Smith, 2011; Khamlichi and Feil, 2018; SanMiguel and Bartolomei, 2018)(REF). Therefore, we measured DNA methylation levels by whole-genome bisulfite sequencing. Reads were assigned to maternal and paternal genomes and DNA methylation levels were calculated from assigned reads per gene. We observed a slightly elevated level of DNA methylation on paternal alleles in wild-type embryos (Fig. S1C). However, we observed a similar paternal bias of DNA methylation in *ez2/ez3* mutants with defective PCR (Fig. S1D-E) and through later stages of development into mature wild-type spores (Fig. S1F-J). Thus, DNA methylation does not appear to be closely associated with the maintenance of PCR.

### Paternal Chromosome Repression is lost by the end of meiosis

To understand when the paternal bias of H3K27me3 causing PCR in Marchantia embryos is lost, we profiled H3K27me3 from various stages of embryo development and determined the ratio of maternal to paternal reads. Two natural accessions from different populations were used to maximize the number of single-nucleotide polymorphisms (SNPs), with Cam-2 as the mother and Tak-1 as the father. We collected embryos at thirteen, sixteen, eighteen and twenty-one days after fertilization (daf) (Figure 1A), sorted nuclei, and profiled chromatin modifications using CUT&RUN (Skene and Henikoff, 2017; Zheng and Gehring, 2019). After assigning H3K27me3 reads to maternal or paternal genomes, we calculated maternal ratios for each gene. At 16 daf, a strong paternal bias of H3K27me3 was observed for 96% of genes, similar to the 88% of paternally biased genes at 13 daf (Figure 1B, S2A-B). However, two days later at 18 daf, the percentage of paternally biased genes decreased to 52% (Fig. 1B, S2C-D; 18 daf). By 21 daf, only 8% of genes showed a paternal H3K27me3 bias (Fig. 1B, S1E-F). There was no correlation of the maternal ratios associated with each gene between the two developmental stages 13 daf and 18 daf nor between the stages 13 daf and 21 daf (Fig. S2G-H), suggesting that genes lost their imprinted status simultaneously. We conclude that paternal imprinting is lost by 21 daf.

To evaluate a potential role for other chromatin marks in PCR, we profiled H2AK119ub deposited by the Polycomb Repressive Complex 1, H3K9me1 that marks constitutive heterochromatin, and H3K36me3 which is associated with transcriptional elongation. In line with our prior observations of H3K9me1 and H3K36me3 at 13 daf (Montgomery et al., 2022), we observed no paternal bias of these marks in any tissue (Fig. 1C). Surprisingly, H2AK119ub was not paternally biased (Fig. 1C) and was thus decoupled from H3K27me3, in contrast with the tight interplay of H2AK119ub and H3K27me3 in non-canonical imprinting in mouse (Mei et al., 2021).

We next aimed to determine the cellular and developmental events that took place prior to the loss of PCR. Diploid Marchantia embryos eventually differentiate spore mother cells, which undergo meiosis to produce tetrads of four connected haploid spores (Brown et al., 2010). The tetrads are separated by a few elongated elater cells that undergo cell death and are contained inside an epithelial cell layer. Hence > 95% of the cells of the embryos are spores (Shimamura, 2015).

These spores undergo maturation, separate from the tetrads and are released at the end of sporogenesis between 26 daf and 28 daf. As embryos develop asynchronously in Marchantia female reproductive organs, we collected, fixed, and imaged individuals from 17 daf to 21 daf (Fig 1D, S2I). At 20 daf, spore tetrads were visible in 50% of individuals (Fig. 1D-E, S2I). At 21 daf, all individuals contained only spore tetrads (Fig. 1D-E) indicating the completion of meiosis and we therefore refer to this stage as post-meiotic spore tetrads (Shimamura, 2015). Thus, the loss of paternal imprinting is completed by the end of meiosis. These findings suggest PCR is confined to the diploid stage of the Marchantia life cycle and that meiosis plays a role in epigenetic reprogramming.

### Reprogramming of the chromatin landscape after meiosis

While one aspect of reprogramming, the erasure of the imprinted state, was completed by the end of meiosis, it remained unclear whether the chromatin landscape of these haploid post-meiotic spore tetrads had been reprogrammed to that of the haploid vegetative state. The vegetative chromatin landscape of Marchantia genes is characterized by H3K36me3 enrichment over gene bodies of transcriptionally active genes and large blocks of H3K27me3 over repressed genes and some transposable elements (Hisanaga et al., 2022; Montgomery et al., 2020). To determine if the non-imprinted post-meiotic chromatin landscape resembled that of haploid vegetative tissue, we compared profiles of chromatin marks in post-meiotic spore tetrads and vegetative tissue. We performed k-means clustering on profiles of H3K27me3, H3K36me3, H3K9me1, and H2AK119ub from embryonic (13 daf), post-meiotic spore tetrads (21 daf) and vegetative tissues (Cam-2) and produced four clusters, labeled A-D (Fig. 2A, S3A). Apart from H3K36me3, the profiles of chromatin marks in post-meiotic spore tetrads did not resemble those of vegetative tissue (Fig. 2A, S3A). Additionally, we called peaks for each chromatin mark and assigned peaks to genes to compare among tissues. There were 1241 genes with peaks of H3K27me3 in vegetative tissue and only seventeen in post-meiotic spore tetrads, of which only four were shared (Fig. 2B), emphasizing the dissimilarity of this mark in these tissues and the erasure of the imprinting marked by H3K27me3. A similar trend was observed for H2AK119ub (Fig. S2B). The dissimilarity of post-meiotic and vegetative chromatin landscapes was also shown in a Pearson correlation matrix of chromatin mark profiles, with only H3K36me3 profiles being moderately correlated, though embryonic H3K36me3 profiles were similarly correlated (Fig. 2C). Additionally, the majority of genes with H3K36me3 peaks in post-meiotic spore tetrads were shared with vegetative tissue (3493 out of 3893; Fig. S3C) reflecting their presence on housekeeping genes (Fig. S3D). Notably, 4388 out of 7980 genes with H3K36me3 peaks in vegetative tissue are unique (Fig. S3C), though this may reflect the diverse set of cell types present in vegetative tissues compared with the uniformity of spore tetrads. Few genes with H3K9me1 peaks overlapped with genes in all tissues, and only four of these were shared among tissues (Fig. S3D). We conclude that the post-meiotic chromatin landscape is unlike that in imprinted embryos, due to the lack of H3K27me3 paternal bias, showing the erasure of the imprinted status. However, post-meiosis, the profiles of chromatin marks are distinct from vegetative tissue, suggesting that meiosis coincides with the erasure of the epigenetic landscape of the embryo, that is required for the *de novo* establishment of the chromatin landscape of the haploid vegetative phase of development.

**Figure 2.**
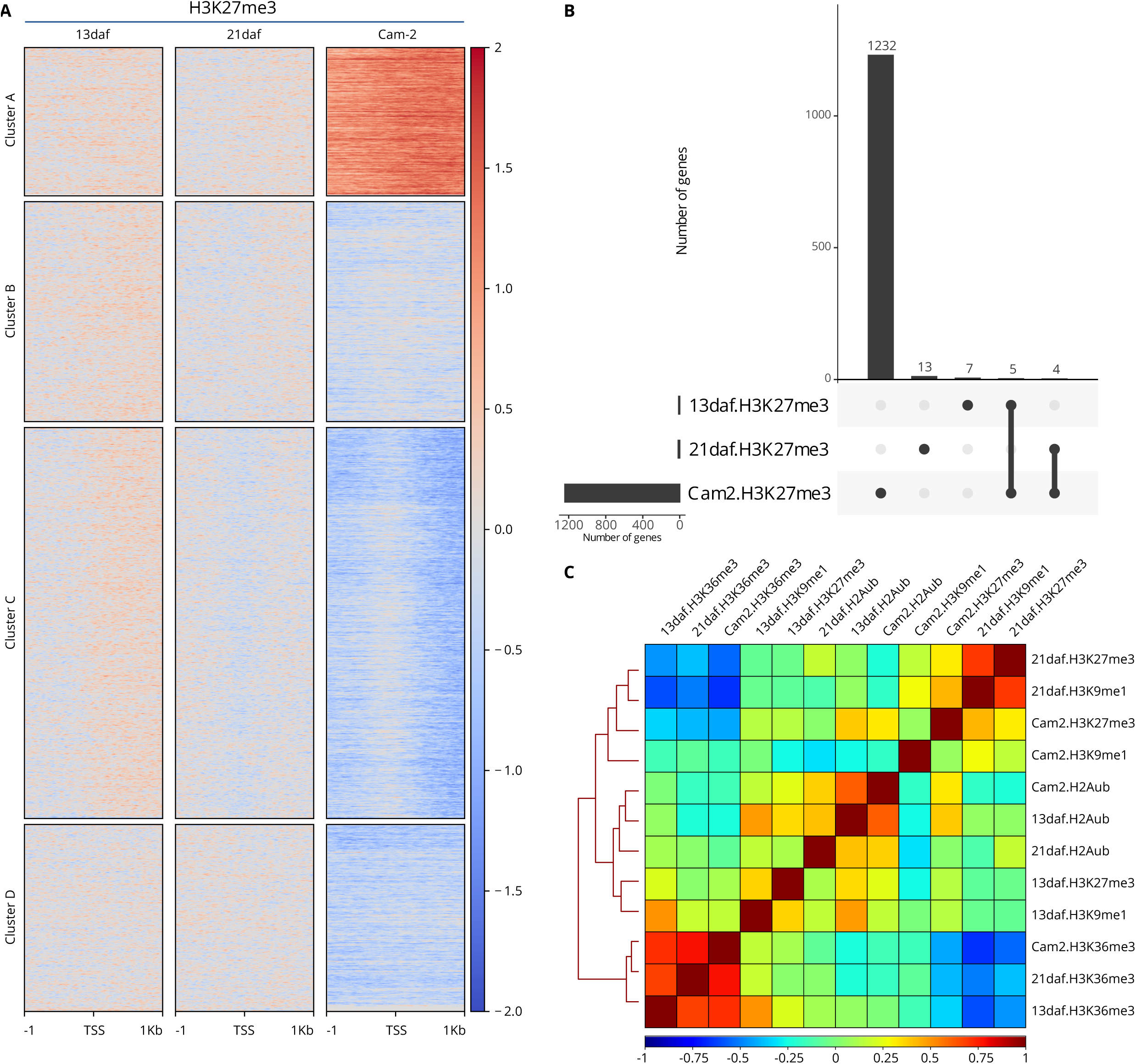
The chromatin landscape in post-meiotic spore tetrads is not reprogrammed to vegetative state. (A) Heatmap of H3K27me3 chromatin profiles centered on gene transcription start sites (TSS) and extending 1 kilobase up- and downstream. K-means clustering was performed using profiles of H3K27me3, H2AK119ub, H3K9me1 and H3K36me3 from embryonic (13 daf), post-meiotic spore tetrad (21 daf) and vegetative (Cam-2) tissue. All profiles are log2 normalized to H3 profiles from the respective tissues. (B) Upset plot of H3K27me3 peaks overlapping genes. Horizontal bars show the total number of genes with H3K27me3 peaks per sample. Vertical bars show the number of shared genes with H3K27me3 peaks among samples for a given comparison, which is shown by filled circles connected by black lines. (C) Pearson correlation matrix of chromatin profiles.

### The vegetative chromatin landscape is established during sporogenesis

As we could not identify the establishment of the vegetative chromatin landscape in post-meiotic spore tetrads (21 daf), we profiled chromatin marks in developing spores (25 daf) and dried, mature spores. K-means clustering of profiles of H3K27me3, H3K36me3, H3K9me1, and H2AK119ub from post-meiotic spore tetrads, developing spores, mature spores, and vegetative tissues produced four clusters, labeled 1-4 (Fig. 3A, S4A). In developing spores, the distribution of each mark over genes already resembled that of mature spores, and vegetative tissue (Fig. 3A, S4A). Likewise, there was a high correlation between developing spores, and vegetative genome-wide profiles of H3K27me3, H2AK119ub and H3K36me3 (Fig. S4B). Further inspection of the four k-means clusters showed that H3K27me3 levels did not differentiate clusters in post-meiotic spore tetrads (Fig. 3B, S4A, 21 daf), whereas Cluster 1 was subsequently characterized by high levels of H3K27me3 in developing spores, mature spores, and vegetative tissues (Fig. 3C-E, S4A). H2AK119ub levels were similarly indistinguishable among post-meiotic spore tetrad clusters (Fig. S4A, S5A), but were elevated in both Clusters 1 and 2 in developing spores, mature spores and vegetative tissue (Fig. S4A, S5B-D). In contrast, Cluster 3 was characterized by elevated levels of H3K36me3 in all tissues (Fig. S5E-H), and by the highest levels of gene expression (Fig. S5I-L). Despite the low level of gene expression and H3K36me3 levels, Cluster 4 did not appear to have elevated levels of H3K27me3 nor H2AK119ub, as in Clusters 1 and 2 (Fig. 3B-E, S5A-L). H3K9me1 enrichment levels were low across all clusters and tissues, with the slight exception of Cluster 1 in post-meiotic spore tetrads (Fig. S5M-P).

**Figure 3.**
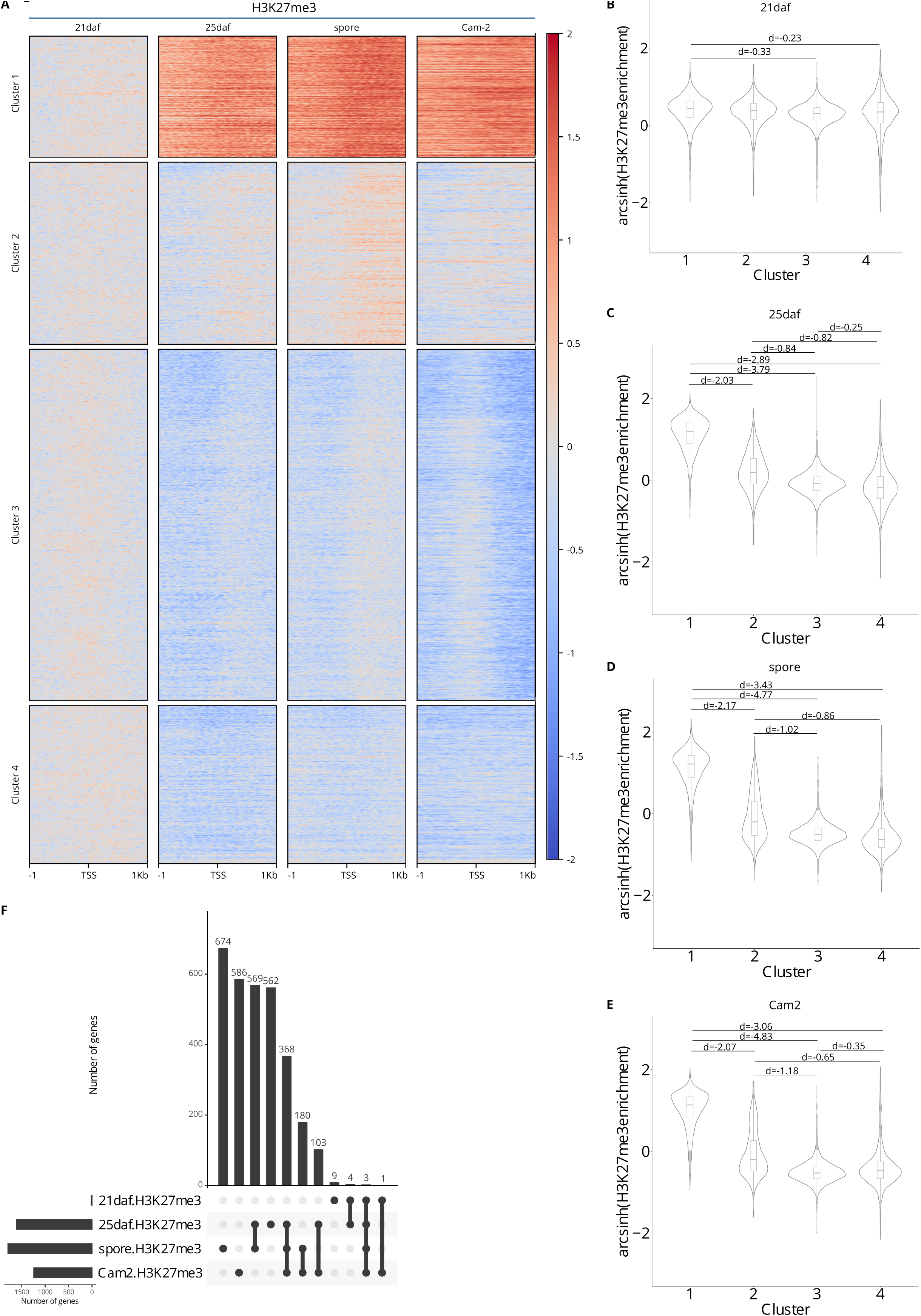
The vegetative chromatin landscape is established in spores. (A) Heatmap of H3K27me3 chromatin profiles centered on gene transcription start sites (TSS) and extending 1 kilobase up- and downstream. K-means clustering was performed using profiles of H3K27me3, H2AK119ub, H3K9me1, and H3K36me3 from post-meiotic spore tetrad (21 daf), developing spore (25 daf), mature spore, and vegetative (Cam-2) tissue. All profiles are log2 normalized to H3 profiles from the respective tissues. (B-E) Violin plots of H3K27me3 enrichment per cluster in (B) post-meiotic spore tetrads (21 daf), (C) developing spore (25daf), (D) mature spores, and (E) vegetative (Cam-2) tissue. Values are arcsinh transformed log2 ratios relative to H3. Non-negligible Cohen’s *d* effect size values are indicated for pairwise comparisons of clusters, where 0.2 < |*d*| < 0.5 is a small effect, 0.5 < |*d*| < 0.8 is a medium effect, and 0.8 < |*d*| is a large effect, as previously reported (Cohen, 1992). (F) Upset plot of H3K27me3 peaks overlapping genes. Horizontal bars show the total number of genes with H3K27me3 peaks per sample. Vertical bars show the number of shared genes with H3K27me3 peaks among samples for a given comparison, which is shown by filled circles connected by black lines.

To further investigate the similarity between chromatin landscapes of developing spores, mature spores, and vegetative tissues, we called peaks for each chromatin mark, assigned them to genes, and compared the overlap of genes among tissues. 53% of the H3K27me3 peaks overlapping genes in vegetative tissue were also found in either or both developing spores and mature spores (Fig. 3F), versus 35% of H2AK119ub peaks (Fig. S5Q). H3K36me3 peaks over genes were highly concordant, with 39% of peaks from vegetative tissue shared in all developmental stages and 34% of vegetative peaks shared among developing spores, mature spores, and vegetative tissues (Fig. S5R). In contrast, H3K9me1 peaks were specific to developmental stages (Fig. S5S). Overall, the vegetative chromatin landscape observed after spore germination and development appears to be established earlier in developing spores within only four days after the completion of meiosis, showing that the completion of epigenetic reprogramming takes place in the products of meiosis.

## Discussion

In conclusion, we found that the restoration of the haploid chromatin landscape involves two steps. First, PCR is reprogrammed by the end of meiosis leading to the erasure of H3K27me3 controlled imprinting. Second, a reprogramming step in the early meiotic spore development reestablishes the chromatin landscape of the vegetative haploid phase prior to spore maturation (Fig. 4).

**Figure 4.**
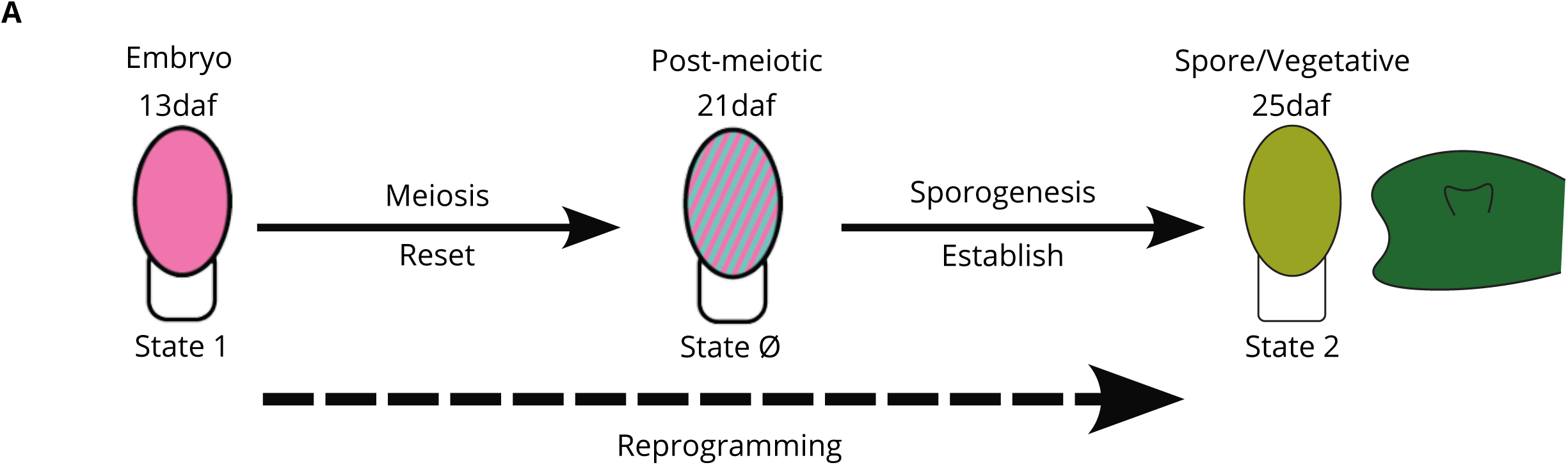
Model of PCR reprogramming in Marchantia. The embryonic chromatin landscape at 13 days after fertilization (daf) (State 1) is characterized by a paternal bias of H3K27me3 over the whole genome. Upon the conclusion of meiosis, the post-meiotic chromatin landscape at 21 daf represents a reset state, with no clear connection between repressive chromatin modifications and gene expression (State Ø). During early spore development, the vegetative chromatin landscape is established (State 2), thus leading to the conclusion of epigenetic reprogramming.

The first step entails the loss of paternally biased H3K27me3, likely completed by the end of meiosis. We were not able to determine whether this occurs via erasure of H3K27me3 from the entire paternal genome, deposition of H3K27me3 over the entire maternal genome, shuffling of paternal nucleosomes marked with H3K27me3 to the maternal genome, or a combination of some or all of these processes. This step is accompanied by a global loss of peaks for other marks, suggesting the involvement of nucleosome exchange. It is possible that, in contrast to mitosis that preserves the chromatin landscape (Flury et al., 2023; Stewart-Morgan et al., 2020), DNA replication at meiosis is followed by *de novo* assembly of nucleosomes without recycling old, modified histones. In addition, meiosis is initiated with numerous DNA double strand breaks (DSBs). Repair of DSBs causes a reshuffling of the chromatin composition over several megabases (García Fernández and Fabre, 2022) In the moss *Physcomitrium patens*, repair of DSBs reprograms the vegetative cell fate (Gu et al., 2020). In plants and animals, hundreds of DSBs initiate meiosis(Choi et al., 2018; Jin et al., 2021). Hence, we predict that DSB repair that initiates meiosis has the potential to reprogram the chromatin landscape in Marchantia.

The second step that completes epigenetic reprogramming is the establishment of the vegetative chromatin landscape after meiosis during sporogenesis. Within four days after meiosis, the chromatin landscape of developing spores largely resembles that of vegetative tissue, thus indicating a completion of the reprogramming process. While the mechanisms involved remain unknown, the absence of mitotic division in developing spores suggests that deposition of the new chromatin landscape is independent of DNA replication and guided by regulators of transcription such as pioneer and general transcription factors.

In mammals, reprogramming of imprinting takes place in the germline (Nashun et al., 2015). Although these steps are thought to be related to germline development rather than to meiosis itself, it was shown that TADs are reconfigured during male meiosis in mammals (Alavattam et al., 2019; Patel et al., 2019; Wang et al., 2019). In flowering plants, chromatin accessibility undergoes a major change during male meiosis (Borg et al., 2021; Nelms and Walbot, 2022). As the mechanisms of meiosis are strongly conserved across eukaryotes (Gerton and Hawley, 2005), the direct impact of chromatin reshuffling during meiosis as a prerequisite to the establishment of totipotency by regulators of transcription is likely a general feature.

## Figure Legends

## Supplementary Figure Legends

**Supplementary Figure 1.**
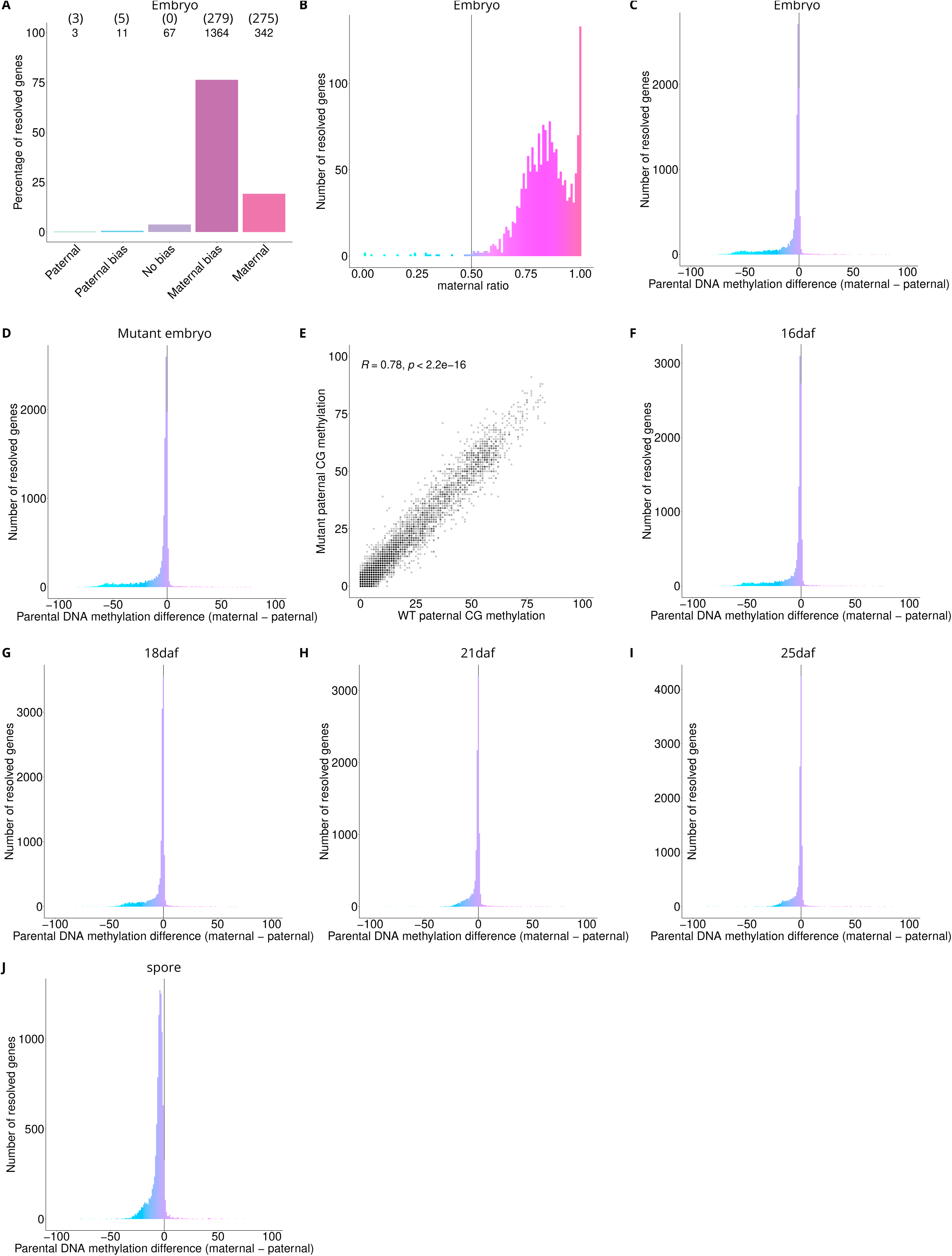
Paternal H3K27me3 bias is lost by 21 daf. (A) Percentage of measured genes within each category of maternal ratio (*p_m_*) of transcription for Cam-2 x Cam-1 embryos. Segments are for paternal (*pm* ≤ 0.05), paternal bias (0.05 < *pm* ≤ 0.35), no bias (0.35 <*pm* < 0.65), maternal bias (0.65 ≤*pm* < 0.95), and maternal (0.95 ≤ *pm*), with the number of genes indicated above each bar and the number of genes significantly deviating from *pm* = 0.5 in parentheses. Significance was assessed using an exact binomial test with Bonferroni correction. (B) Histogram of the maternal ratio (*pm*) of transcription per gene for Cam-2 x Cam-1 embryos. Each bin is 0.01 units wide. (C) Histogram of the difference of CG DNA methylation percentage between maternal and paternal reads per gene for wild-type Cam-2 x Tak-1 embryos. (D) As (C) for *ez2/ez3* mutant embryos. (E) Scatterplot of wild-type embryo paternal CG methylation versus *ez2/ez3* mutant embryo paternal CG methylation. Spearman correlation is indicated. (F) As (C) for 16 daf. (G) As (C) for 18 daf. (H) As (C) for 21 daf. (I) As (C) for 25 daf. (J) As (C) for spores.

**Supplementary Figure 2.**
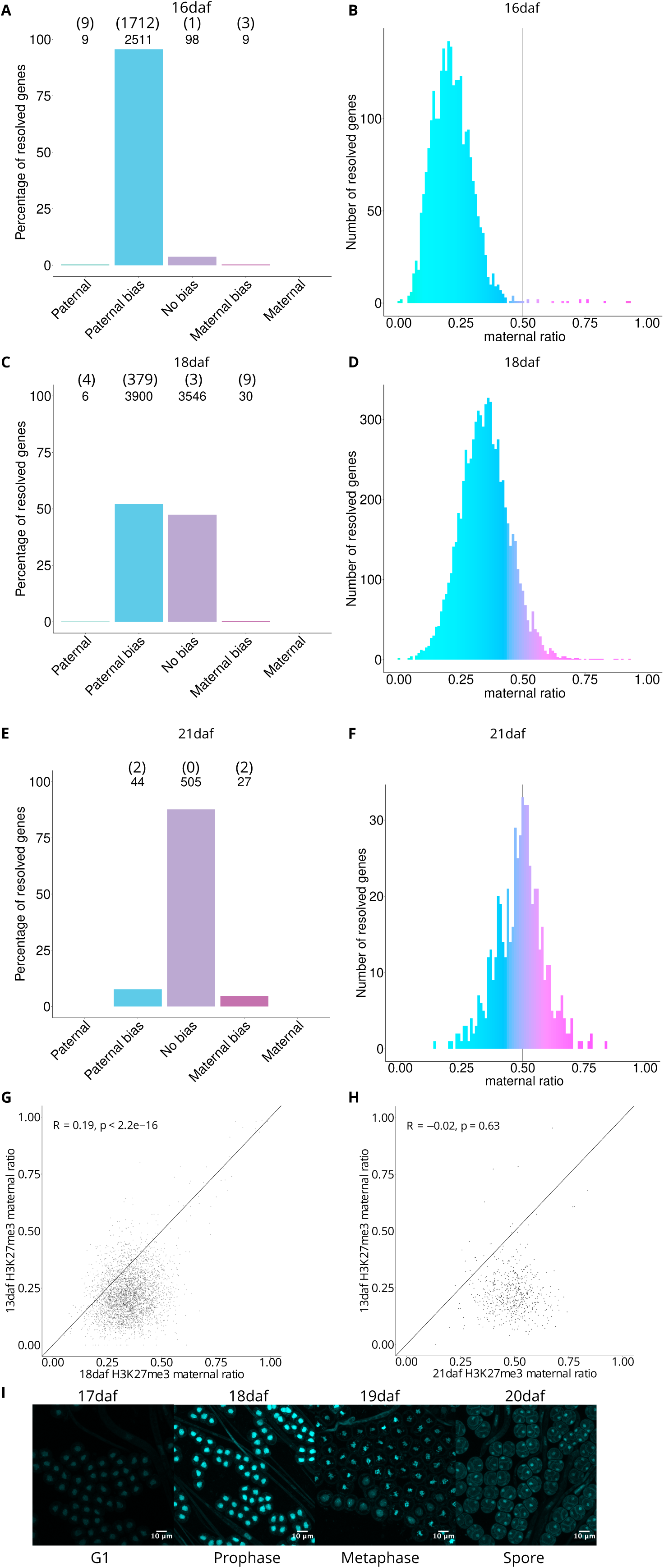
Paternal H3K27me3 bias is lost by 21 daf. (A) Percentage of measured genes within each category of maternal ratio (*p_m_*) of H3K27me3 at 16 days after fertilization (daf). Segments are for paternal (*pm* ≤ 0.05), paternal bias (0.05 < *pm* ≤ 0.35), no bias (0.35 <*pm* < 0.65), maternal bias (0.65 ≤*pm* < 0.95), and maternal (0.95 ≤ *pm*), with the number of genes indicated above each bar and the number of genes significantly deviating from *pm* = 0.5 in parentheses. Significance was assessed using an exact binomial test with Bonferroni correction. (B) Histogram of the maternal ratio (*pm*) of H3K27me3 per gene at 16 daf. Each bin is 0.01 units wide. (C) As in (A) for 18 daf. (D) As in (B) for 18 daf). (E) As in (A) for 21 daf. (F) As in (B) for 21 daf. (G) Scatterplot of 18 daf versus 13 daf H3K27me3 maternal ratio per resolved gene. Pearson correlation is indicated. (H) As in (G) for 21 daf versus 13 daf. (I) Representative DAPI staining of 17 daf, 18 daf, 19 daf, and 20 daf samples.

**Supplementary Figure 3.**
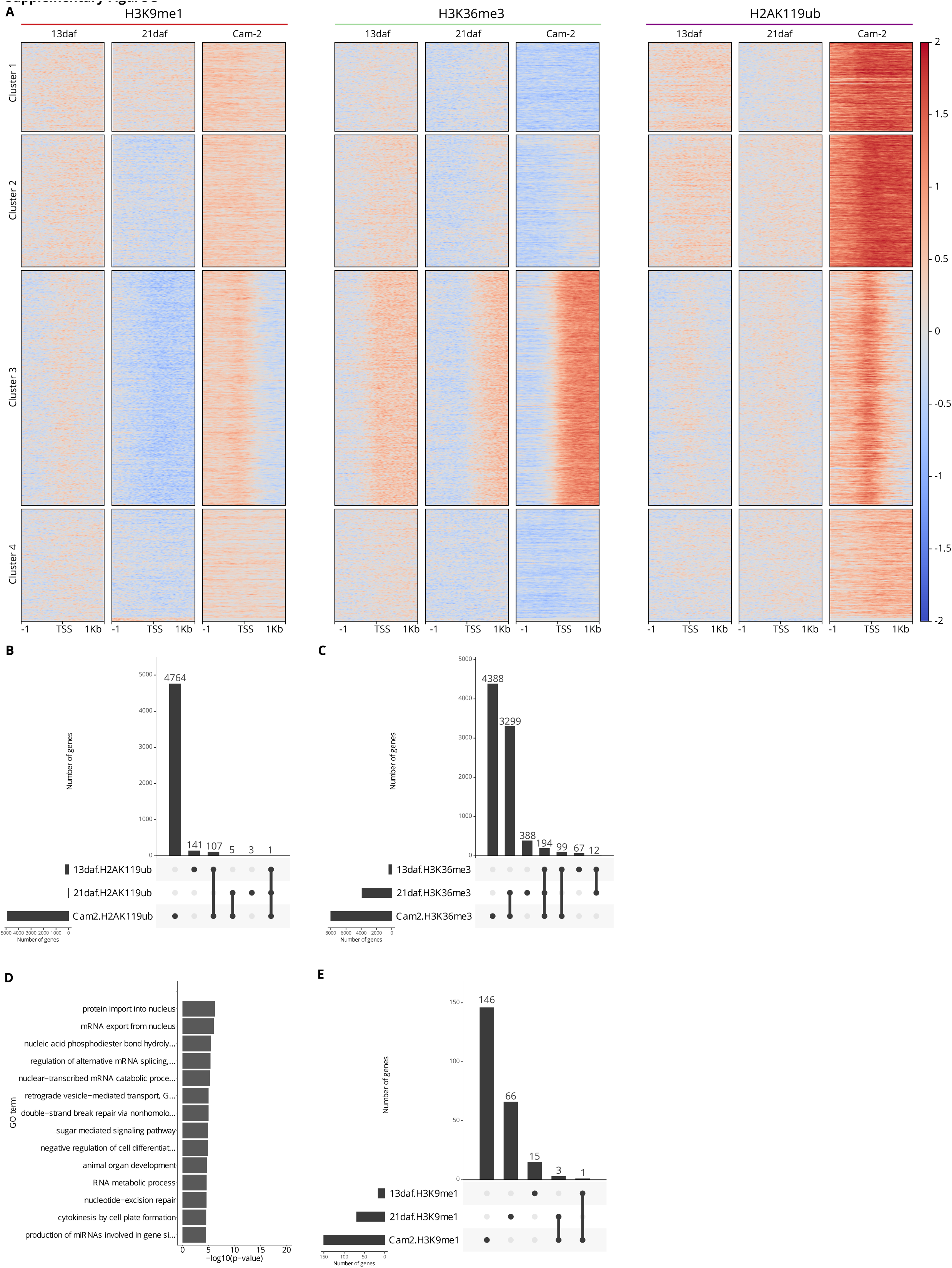
Comparison of additional chromatin marks among embryonic, post-meiotic spore tetrad, and vegetative tissues. (A) Heatmap of H3K9me1, H2AK119ub and H3K36me3 chromatin profiles centered on gene transcription start sites (TSS) and extending 1 kilobase up- and downstream. K-means clustering was performed including profiles of H3K27me3 from embryonic (13 daf), post-meiotic spore tetrad (21 daf) and vegetative (Cam-2) tissue. All profiles are log2 normalized to H3 profiles from the respective tissues. (B-C) Upset plots of (B) H2AK119ub, (C) H3K36me3 peaks overlapping genes. Horizontal bars show the total number of genes with peaks per sample. Vertical bars show the number of shared genes with peaks among samples for a given comparison, which is shown by filled circles connected by black lines. (D) Gene Ontology term enrichment for Biological Processes of genes with H3K36me3 peaks shared between post-meiotic spore tetrads and vegetative tissue. Terms are ordered by p-value. (E) Upset plot of H3K9me1 peaks overlapping genes.

**Supplementary Figure 4.**
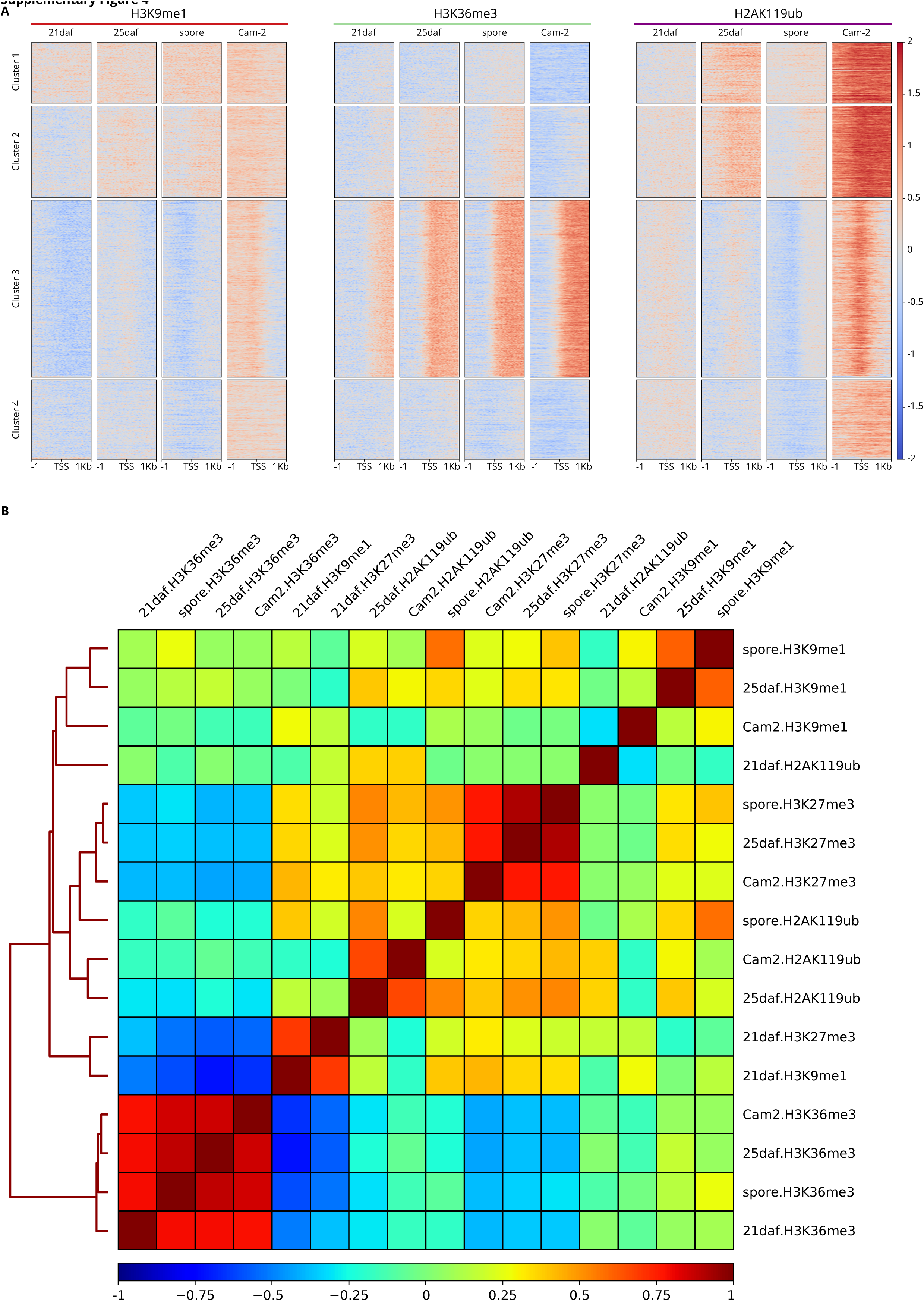
Comparison of additional chromatin marks among post-meiotic spore tetrad, developing spore, mature spore, and vegetative tissues. (A) Heatmap of H3K9me1, H2AK119ub, and H3K36me3 chromatin profiles centered on gene transcription start sites (TSS) and extending 1 kilobase up- and downstream. K-means clustering was performed including profiles of H3K27me3 from post-meiotic spore tetrad (21 daf), developing spore (25 daf), mature spore, and vegetative (Cam-2) tissue. All profiles are log2 normalized to H3 profiles from the respective tissues. (B) Pearson correlation matrix of chromatin profiles.

**Supplementary Figure 5.**
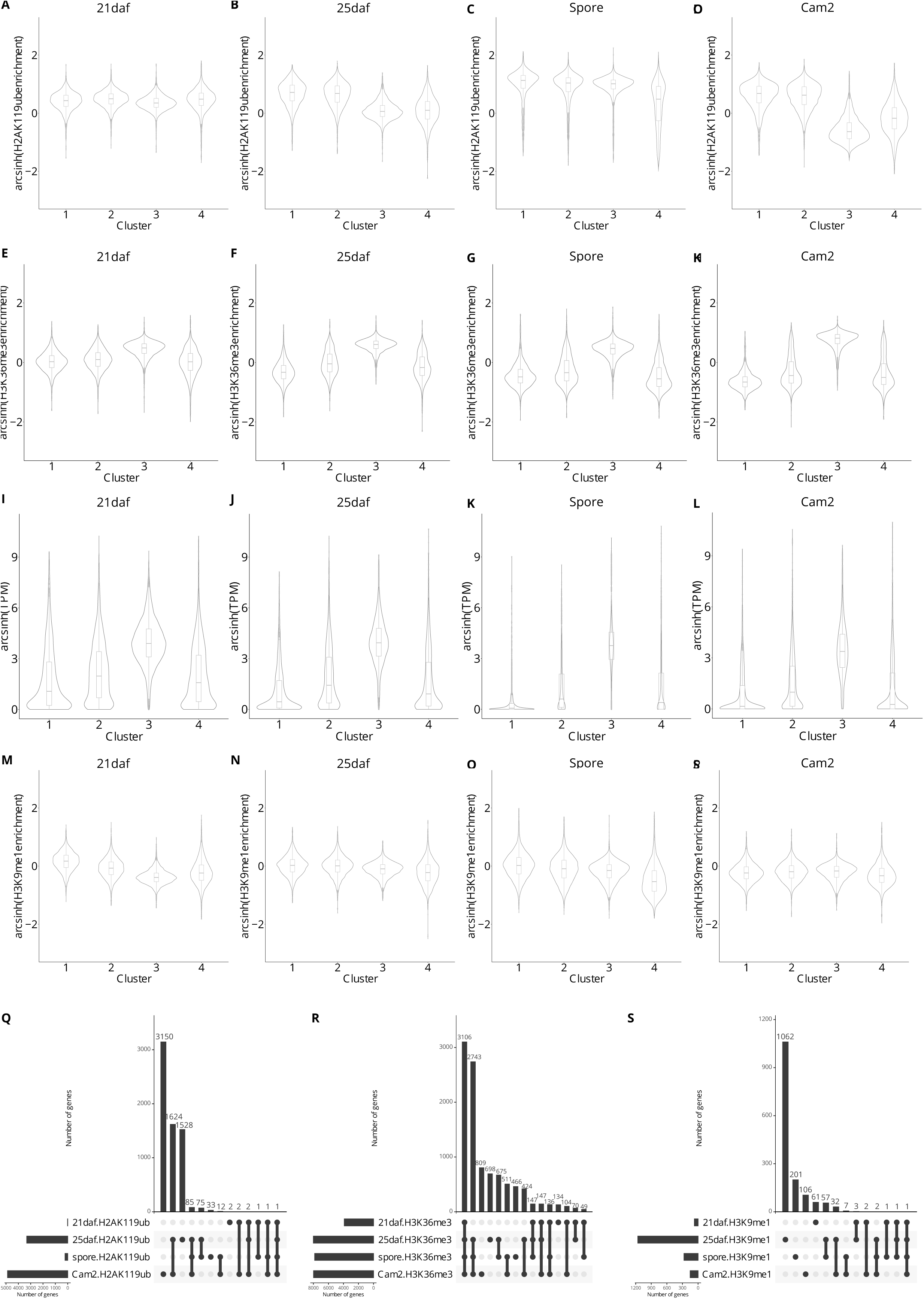
Distributions of chromatin marks over clusters and peaks. (A-D) Violin plots of H2AK119ub enrichment in (A) post-meiotic spore tetrad (21 daf), (B) developing spore (25 daf), (C) mature spore and (D) vegetative tissue per cluster. Values are arcsinh transformed log2 ratios relative to H3. (E-H) Violin plots of H3K36me3 enrichment in (E) post-meiotic spore tetrad (21 daf), (F) developing spore (25 daf), (G) mature spore and (H) vegetative tissue per cluster. Values are arcsinh transformed log2 ratios relative to H3. (I-L) Violin plots of transcripts per million (TPM) in (I) post-meiotic spore tetrad (21 daf), (J) developing spore (25 daf), (K) mature spore, and (L) vegetative tissue per cluster. Values are arcsinh transformed. (M-P) Violin plots of H3K9me1 enrichment in (M) post-meiotic spore tetrad (21 daf), (N) developing spore (25 daf), (O) mature spore, and (P) vegetative tissue per cluster. Values are arcsinh transformed log2 ratios relative to H3. (Q-S) Upset plots of (Q) H2AK119ub, (R) H3K36me3, (S) and H3K9me1 peaks overlapping genes. Horizontal bars show the total number of genes with peaks per sample. Vertical bars show the number of shared genes with peaks among samples for a given comparison, which is shown by filled circles connected by black lines.

## Materials and Methods

### Plant lines and growth conditions

Wild-type male Tak-1 and female Cam-2 accessions of *Marchantia polymorpha* ssp. *ruderalis* were used in this study.

Female plants for crosses were grown at room temperature on Grodan Vital (Grodan, Roermond, The Netherlands) supplemented with liquid Hyponex fertilizer (Hyponex, Osaka, Japan) under constant white light. Male plants for crosses were grown at 22 °C on Neuhaus N3 substrate soil (Humko, Podnart, Slovenia) under 16 hours of far-red light and 80% humidity. Plants grown for the collection or observation of vegetative tissues were grown under axenic conditions on half-strength Gamborg B5 media without vitamins (Duchefa Biochemie, Haarlem, The Netherlands) and 1% (w/v) agar under constant white light.

Crosses were performed by collecting mature antheridiophores in water and pipetting the water containing released sperm onto archegoniophores.

### Transcriptome generation

Sporophyte samples were collected by hand dissection, with one sporophyte per replicate. Sporophytes and the surrounding maternal calyptra tissue were dissected out of the archegoniophores into 10% RNALater (Qiagen, Hilden, Germany) on Microscope slides with cavities (Marienfeld Superior, Lauda-Königshofen, Germany) and the sporophyte was further dissected out of the surrounding maternal tissue (Montgomery et al., 2022). Each embryo was washed four times in a series of wells containing 150 µL 10% RNALater to remove any maternal RNAs, as previously described for the pure isolation of plant embryos (Kao and Nodine, 2020) (Kao and Nodine, 2020). Each embryo was then placed in 30 µL 100% RNALater on ice until sample collection was completed. The solution was diluted to 10% RNALater by the addition of 270 µL RNase-free water (Zymo Research, Irvine, CA), vortexed gently, and the solution removed. Samples were either resuspended in 30 µL 100% RNALater and stored at -70°C or resuspended in 100 µL TRI reagent (Zymo Research). Samples were crushed with a micropestle and RNA was extracted using a Direct-zol RNA MicroPrep kit (Zymo Research).

Spore transcriptomes were generated from three frozen mature sporophytes per replicate. Frozen sporophytes were disrupted in 100 µL DNase/RNase-free water with a pipette tip to release spores, and centrifuged at 13,000 x g for 1 minute to pellet spores. 250 µL of TRI reagent (Zymo Research) was added to each sample plus 50 µL of 0.2 µm glass beads and vortexed for 3 minutes to break spore walls. 250 µL of the sample was transferred to a fresh tube and RNA was extracted using a Direct-zol RNA microPrep kit (Zymo Research). All RNA samples were treated to remove DNA using a DNase Treatment and Removal kit (Invitrogen, Thermo Fisher Scientific Inc, Waltham, MA, USA).

RNA-seq libraries were generated from total RNA following the Smart-seq2 protocol (Picelli et al., 2014) (Picelli et al., 2014). cDNA synthesis was performed on 1 µL of total RNA. 1 µL of 10 µM 5’-Bio-anchored oligo dT ([Btn]AAGCAGTGGTATCAACGCAGAGTACTTTTTTTTTTTTTTTTTTTTTTTTTTTTTTVN) and 1 µL 10mM dNTPs were added to each sample and incubated at 72°C for 3 minutes and immediately placed on ice. 7 µL of a mastermix containing 0.5 µL SuperScript IV (Invitrogen), 0.25 µL RiboLock (ThermoFisher Scientific), 2 µL Superscript IV buffer (Invitrogen), 1 µL 100mM DTT (Invitrogen), 2 µL 5M Betaine (Sigma-Aldrich), 0.9 µL MgCl_2_, 0.1 µL nuclease-free water (Invitrogen), and 1 µL 10 µM 5’-Bio TSO ([Btn]AAGCAGTGGTATCAACGCAGAGTACrGrG+G, Exiqon) was added to each sample and the cDNA synthesis reaction took place under the following thermocycling conditions: 42°C 90 min, <50°C 2 min, 42°C 2 min>x10, 70°C 15 min. 40 µL of a mastermix containing 14.5 µL nuclease-free water, 25 µL Q5 Hot Start 2x MasterMix (New England Biolabs, Ipswich, MA), and 0.5 µL 10 µM 5’ Bio-ISPCR oligo ([Btn]AAGCAGTGGTATCAACGCAGAGT) was added to each sample and the PCR pre-amplification was performed under the following thermocycling conditions: 98°C 3 min, <98°C 15 s, 67°C 20 s, 72°C 6 min>x12, 72°C 5 minutes. Samples were cleaned up by bead purification using 1x volume of MBSPure beads (Molecular Biology Services, IMP, Vienna, Austria) and eluted in 15 µL of 10mM Tris-HCl. 5-50 ng of each sample was used for the tagmentation reaction, containing 2.5 µL of 4x TAPS-DMF buffer and 1 µL of Tn5 (Molecular Biology Services, IMP), which was performed for 3 minutes at 55°C, after which samples were immediately placed on ice. Samples were purified using a DNA Clean and Concentrator kit (Zymo Research) using manufacturer’s instructions and eluted in 10 µL of 10mM Tris-HCl. Tagmented samples were amplified by the addition of 2.5 µL each of 10 µM barcoded forward and reverse primers (Picelli et al., 2014) (Picelli et al., 2014) and 15 µL Q5 2x HiFI MasterMix (New England Biolabs) using the following thermocycling conditions: 72°C 3 min, 98°C 20 s, <98°C 10 s, 63°C 30 s, 72°C 3 min>x5. Amplified samples were cleaned up by bead purification using 1x volume of MBSPure beads (Molecular Biology Services, IMP). Samples were sequenced on an Illumnia NovaSeq to generate 50bp paired-end reads. Nine biological replicates of 21 daf, nine of 25 daf and three of spores were used for subsequent analyses.

### Chromatin profiling by CUT&RUN

Sporophytes at 16, 18, 21 and 25 days after fertilization and the surrounding calyptra were hand-dissected from archegoniophores. All sporophyte samples were collected into Galbraith buffer (45 mM MgCl_2_-6H2O, 30 mM Trisodium citrate, 20 mM MOPS) pH 7.0 plus 0.1% Triton X-100 and 1x cOmplete Protease Inhibitor Cocktail (Roche, Mannheim, Germany) on ice. Samples were crushed using a mortar and pestle on ice to release nuclei. 21 daf sporophytes were incubated with 1% pectolyase, 1% cellulase and 1% beta-glucuronidase (Duchefa Biochemie, Haarlem, The Netherlands) for 10 minutes at 37 °C. 250uL aliquots of 21 daf and 25 daf samples were transferred to 1.5mL tubes containing 50 µL 0.2 µM glass beads and vortexed for 3 minutes. All samples were then filtered through a 40 µm filter (VWR, Radnor, PA, USA) before staining with 2 µg/mL DAPI. For spore samples, 14 aliquots of ten mature sporophytes each were collected in 1.5mL tubes and frozen at -70°C. Spores were released by manual disruption of frozen sporophytes in 100 µL of water and aliquots were combined in two 5mL tubes. Spores were pelleted by spinning at 13,000 x g for 1 minute, the supernatant was removed and the pellet was resuspended in 250 µL of Galbraith buffer pH 7.0 plus 0.1% Triton X-100 and 1x cOmplete Protease Inhibitor Cocktail (Roche, Mannheim, Germany). 50 µL of 0.2 µm glass beads were added and samples were vortexed for 3 minutes to smash open spores and release nuclei. The nuclei suspension was transferred to a new tube, filtered through a 40 µm filter and stained with 2 µg/mL DAPI. Cam-2 gemmae were grown under axenic conditions on half-strength Gamborg B5 media without vitamins (Duchefa Biochemie, Haarlem, The Netherlands) and 1% (w/v) agar under constant white light for 14 days. 1 g of green thallus was harvested in Galbraith buffer pH 7.0 plus 0.1% Triton X-100 and 1x cOmplete Protease Inhibitor Cocktail (Roche, Mannheim, Germany) on ice. Samples were chopped with a razor blade in a Petri dish on ice to release nuclei and were filtered through a 40 µm filter before staining with 2 µg/mL DAPI.

Nuclei were sorted on a BD FACSARIA III (BD Biosciences, San Jose, CA, USA). 2n nuclei were sorted for 16 daf and 4n nuclei were sorted for 18 daf to deplete contaminating maternal tissue not removed during sample harvesting. 1n nuclei were sorted for 21 daf, 25 daf, spore, and Cam-2 samples. Samples were sorted into 100 µL of Wash buffer (20 mM HEPES pH 7.5, 150 mM NaCl, 0.5 mM Spermidine, 1x cOmplete Protease Inhibitor Cocktail (Roche)) with 40,000 nuclei per replicate. Bio-Mag Plus Concanavalin A coated beads (Polysciences, Inc., Warrington, PA, USA) were activated by mixing 10 µL ConA beads per sample in 1.5mL Binding buffer (20 mM HEPES-KOH pH 7.9, 10 mM KCl, 1 mM CaCl_2_, 1 mM MnCl_2_). The beads were placed on a magnet, liquid was removed, and the beads were resuspended in 1.5 mL Binding buffer. Liquid was again removed from the beads on a magnet and beads were resuspended in 10 µL Binding buffer per sample. 10 µL of the activated beads were added to each sorted nuclei sample and incubated at room temperature for 10 min on a rotator. Liquid was removed from the bead-bound nuclei on a magnet and samples were resuspended in 50 µL Antibody buffer (Wash buffer plus 2 mM EDTA). 0.5 µL of each antibody (H3K27me3 Millipore, Temecula, CA, USA, #07-449 RRID:AB_310624; H3K36me3 Abcam, Cambridge, UK, ab9050 RRID:AB_306966; H3K9me1 Abcam ab9045 RRID:AB_306963; H2AK119ub Cell Signaling Technology D27C4 RRID:AB_10891618; H3 Abcam ab1791 RRID:AB_302613) used was added to samples while gently vortexing and samples were incubated overnight at 4°C on a shaker. Liquid was removed from the samples on a magnet and washed twice in 1 mL Wash buffer before resuspending in 50 µL Wash buffer. 1.16 µL of 30 µg/mL pAG-MNase (Molecular Biology Service, IMP) was added to each sample by gently vortexing and placed on a shaker for 10 min at room temperature. Liquid was removed from the samples on a magnet and washed twice in 1 mL Wash buffer before resuspending in 150 µL Wash buffer. 3 µL 100 mM CaCl_2_ was added to ice-cold samples while gently vortexing and shaken at 4°C for two hours. 100 µL STOP buffer (340 mM NaCl, 20 mM EDTA, 4 mM EGTA, 50 µg/mL RNase A (ThermoFisher Scientific), 50 µg/mL glycogen, 10 pg/mL heterologous HEK293 DNA) was added to stop the reaction. Samples were incubated at 37°C for 10 min at 500 RPM then spun at 4°C for 5 min at 16,000 x g. Samples were placed on a magnet and the liquid containing released DNA fragments was transferred to a new tube. 2.5 µL 10% SDS and 2.5 µL 20 mg/mL Proteinase K (ThermoFisher Scientific) was added to each sample, mixed by inversion, and incubated for 1hr at 50°C. 250 µL buffered phenol-chloroform-isoamyl solution (25:24:1) was added to each sample, followed by vortexing and transfer to MaXtract tubes (Qiagen, Hilden, Germany). Samples were spun for 5 min at 16,000 x g. 250 µL chloroform was added and samples were spun for 5 min at 16,000 x g. The top aqueous phase was transferred to a fresh tube containing 2 µL 2 mg/mL glycogen. 625 µL 100% EtOH was added before vortexing and chilling at -20°C overnight. DNA extraction continued with spinning for 10 min at 4°C at 20,000 x g. The supernatant was poured off and 1 mL 100% EtOH was added to the samples before spinning again for 1 min at 4°C at 16,000 x g. Supernatant was discarded and samples air-dried before dissolving in 50 µL 0.1x TE. A NEBNext Ultra II DNA library prep kit for Illumina (New England Biolabs) was used according to the manufacturer’s instructions for sample library preparation. Samples were sequenced on either an Illumina HiSeqv4 or NovaSeq to generate 50bp paired-end reads. At least two biological replicates were used for each sample for H3K27me3, H3K36me3, H3K9me1, H2AK119ub and H3 in 16 daf, 18 daf, 21 daf, 25 daf, spores, and Cam-2.

### Whole-genome bisulfite sequencing by tagmentation

DNA was extracted from thallus samples for whole-genome bisulfite sequencing as follows. 0.1 g of 14 day Cam-2 or Tak-1 thallus per replicate was collected in a Precellys tube (Bertin Instruments, Montigny-le-Bretonneux, France) with five 2.8 mm stainless steel beads (Bertin Corp., Rockville, MD) in liquid nitrogen and disrupted with a Precellys Evolution tissue homogenizer (Bertin Instruments) cooled using liquid nitrogen with the following settings: 4500 RPM 30 seconds, 5 second pause, repeated twice. A PhytoPure DNA Extraction kit (GE Healthcare) was used to complete the DNA extraction. 600 µL Reagent 1 was added and mixed. 200 µL Reagent 2 was added and tubes were inverted to mix. Samples were incubated at 65°C for 10 minutes while shaking before transferring the sample to a fresh 1.5mL tube and incubated on ice for 20 minutes. Samples were removed from ice and 500 µL -20°C chloroform was added. 100 µL Nucleon PhytoPure DNA Extraction Resin was added and samples were placed on a shaker at room temperature for 10 minutes. Samples were centrifuged at 1300 x g for 10 minutes and the upper layer was transferred to a new tube. An equal volume of cold isopropanol was added and tubes were inverted until DNA precipitated. Samples were centrifuged at 4000 x g for 5 minutes. After removing the supernatant, the DNA pellet was washed with cold 70% ethanol before discarding the supernatant and air dried for 10 minutes. The DNA pellet was resuspended in 20 µL 10 mM Tris pH 8 and further cleanup was done with a Zymo DNA Clean and Concentrator kit (Zymo Research).

DNA was extracted from sporophyte samples for whole-genome bisulfite sequencing as follows using a QiAmp Micro DNA Extraction kit (Qiagen, Hilden, Germany). 20-50 sporophytes were collected per replicate depending on the size of each sample, around 50 for 13 daf, 45 for 16 daf, 40 for 18 daf and 25 for 21 daf, and 25 daf. Pure sporophytes dissected out of calyptras were dissected in DNase/RNase-free water in glass well slides and kept on ice in water. Water was removed and 180 µL Buffer ATL was added before warming samples to room temperature. 20 µL Proteinase K was added and samples were pulse-vortexed for 15 seconds and incubated at 56°C overnight. 200 µL Buffer AL was added and samples were pulse-vortexed for 15 seconds. 200 µL 100% ethanol was added and samples were pulse-vortexed for 15 seconds. Samples were incubated at room temperature for 5 minutes before being spun down briefly and transferred to a QIAmp MinElute column. Samples were centrifuged at 6000 x g for 1 minute and columns were moved to a new collection tube. 500 µL Buffer AW1 was added and samples were centrifuged at 6000 x g for 1 minute and columns were moved to a new collection tube. 500 µL Buffer AW2 was added and samples were centrifuged at 6000 x g for 1 minute and columns were moved to a new collection tube. Samples were centrifuged at 20,000 x g for 3 minutes and transferred to a new 1.5mL tube. 30 µL Buffer AE was added to columns, incubated for 1 minute and centrifuged at 20,000 x g for 1 minute.

A tagmentation-based approach was used as in (Wang et al., 2013). The Tn5mC transposase was assembled as follows. Equal amounts of 100 µM primer Tn5mC-Apt1 (TcGTcGGcAGcGTcAGATGTGTATAAGAGAcAG; where c indicates 5C methylated) and 100 µM primer Tn5mC1.1-A1block (pCTGTCTCTTATACAddC; where p indicates phosphate and ddC indicates dideoxycytidylate) were mixed in a PCR tube and thermocycled under the following conditions: 95°C 3 min, 70°C 3 min, 70°C to 25°C for 30 seconds at each step in intervals of 1°C decreases. The annealed oligos were mixed with Tn5 (Molecular Biology Services, IMP, Vienna, Austria) and dilution buffer in a 1.43:1:7.57 ratio, incubated at room temperature for 1 hour, and stored at -20°C until use. Tagmentation was done as follows: 3 µL extracted DNA was mixed with 2 µL 5x TAPS-DMF buffer, 1 µL Tn5mC and 4 µL water and incubated at 55°C for 3 minutes. 7.5 µL 5M guanidinium thiocyanate was added and mixed by pipetting. Samples were cleaned up with a Zymo DNA Clean and Concentrator kit (Zymo Research) and eluted in 12 µL EB. Oligo replacement and gap repair was done as follows: 11 µL sample was mixed with 2 µL 10 mM dNTP, 2 µL 10x Ampligase buffer and 2 µL 10 µM Tn5mC-Repl01 oligo (pcTGTcTcTTATAcAcATcTccGAGccCAcGAGAcinvT; where p indicates phosphate, c indicates 5C methylated and invT indicated inverted deoxythymidylate) and thermocycled under the following conditions: 50°C 1 minute, 45°C 10 minutes followed by a decrease of 0.1°C per second down to 37°C. While samples remained in the thermocycler, 1 µL T4 DNA polymerase (New England Biolabs, Ipswich, MA) and 2.5 µL Ampligase (Biozym) were added and samples were mixed by pipetting before incubating for an additional 30 minutes at 37°C. Samples were cleaned up by bead purification using 1.3x volume of MBSPure beads (Molecular Biology Services, IMP, Vienna, Austria) and samples were eluted in 22 µL water, with 2 µL of sample set aside to make non-bisulfite-treated libraries. Bisulfite treatment was performed with a EZ DNA Methylation-Gold kit (Zymo Research) as follows: CT conversion reagent was warmed to 37°C and 130 µL was added to 20 µL sample. Samples were pipetted to mix, spun down briefly and split into two 75 µL aliquots in PCR tubes. Thermocycling was done under the following conditions: 98°C 10 minutes, 64°C 2.5 hours, 4°C for less than 20 hours. 600 µL M-binding buffer was added to columns and the sample was then added and inverted to mix before centrifuging at 11,000 x g for 30 seconds. 100 µL M-Wash buffer was added to the column and centrifuged for 30 seconds. 200 µL M-desulphonation buffer was added and incubated at room temperature for 15 minutes before centrifuging for 30 seconds. The columns were washed twice with 200 µL M-Wash buffer with 30 second centrifuges. Sample were centrifuged at 17,000 x g for 3 minutes and columns were transferred to Lo-Bind tubes. 12 µL M-elution buffer was added and samples were spun at 16,000 x g for 30 seconds. A second elution in the same tube with 12 µL M-elution buffer was done. Libraries were amplified as follows. 22 µL sample was mixed with 25 µL 2x KAPA 2G robust HS Master Mix (Sigma-Aldrich, Merck KGaA, Darmstadt, Germany), 1.5 µL 10 µM Index 1 primer and 1.5 µL 10 µM Index 2 primer. Thermocycling as follows: 95°C 3 minutes, (95°C 20 seconds, 62°C 15 seconds, 72°C 40 seconds)x7. A 2.5 µL aliquot of the samples was taken for qPCR to check the number of cycles using the NEB Luna 2x mix (New England Biolabs, Ipswich, MA). After completing the required number of additional PCR cycles, samples were cleaned up by bead purification using 1.8x volume of MBSPure beads (Molecular Biology Services, IMP, Vienna, Austria) and samples were eluted in 32 µL EB. Samples were sequenced on an Illumnia NovaSeq to generate 50bp paired-end reads.

### Meiosis imaging

Eight individual sporophytes were collected from Cam-2 x Tak-1 crosses at 17, 18, 19, 20, and 21 days after fertilization (daf). Samples were collected in 100 µL PME buffer (50 mM Pipes, 5 mM EGTA, 1 mM MgSO_4_ 7H_2_O) plus 4% formaldehyde and 0.3 M Mannitol per sporophyte on ice and fixed for 30 minutes at room temperature under a vacuum. Samples were washed three times with PME buffer for 10 minutes on ice and mounted on glass slides in PME buffer plus 5 µg/L Hoechst 33342 (ThermoFisher Scientific) and squashed with a cover slip before sealing.

Images were acquired with an LSM 780 scanning laser confocal microscope (Zeiss).

### Transcriptome analysis

Published transcriptomes from wild-type 13 day after fertilization Cam-2 x Tak-1 embryos and Cam-2 and Tak-1 vegetative tissues (Montgomery et al., 2022) were downloaded from the SRA database.

Reads were mapped to the Takv6 genome (Iwasaki et al., 2021) wherein all SNP positions between Tak-1 and Cam-2 were replaced with N’s, where SNPs were identified in (Montgomery et al., 2022). Reads were preprocessed with SAMtools v1.9 (Li et al., 2009) and BEDTools v2.27.1 (Quinlan and Hall, 2010), trimmed with Trim Galore (https://github.com/FelixKrueger/TrimGalore) and mapped with STAR v2.7.1 (Dobin et al., 2013). Transcripts per Million (TPM) values were calculated by RSEM v1.3.2 (Li and Dewey, 2011). Data from RSEM were imported into R v3.5.1 (R Core Team, 2018) using the tximport package v1.10.1 (Soneson et al., 2015). Differential gene analysis was performed using DeSeq2 v1.22.2 (Love et al., 2014). Principal component analysis was performed in R v3.5.1 (R Core Team, 2018). Heatmaps were generated in R using the pheatmap v1.0.12 package (Kolde, 2019).

### CUT&RUN data analysis

Reads were mapped to the Takv6 genome (Iwasaki et al., 2021) wherein all SNP positions between Tak-1 and Cam-2 were replaced with N’s, where SNPs were identified in (Montgomery et al., 2022). File processing and mapping parameters were performed as previously published (Montgomery et al., 2020). Chromatin enrichment per gene was calculated by counting the number of reads and normalizing to 1x genome coverage. Upset plots were generated in R using the UpSetR v1.4.0 package (Conway et al., 2017). K-means clustering of chromatin mark profile heatmaps were performed and visualized with deepTools v3.3.1 (Ramírez et al., 2016), as was hierarchical clustering of Pearson correlations of chromatin mark samples. Plots were made in R using the ggplot2 package v3.3.2 (Wickham, 2016) and Pearson correlations were calculated with the ggpubr v0.4.0 package (Kassambara, 2020). Effect size (Cohen’s *d*) was calculated in R using effsize v0.7.6(Torchiano, 2020) where |d| < 0.2 is no effect, 0.2 < |d| < 0.5 is a small effect, 0.5 < |d| < 0.8 is a medium effect, and |d| > 0.8 is a large effect, as previously reported (Cohen, 1992).

Chromatin mark peaks were called using MACS2 v2.2.5 (Feng et al., 2012) and considered to associate with a gene if it overlapped with at least 50% of the peak length using BEDTools v2.27.1 (Quinlan and Hall, 2010).

### DNA methylation data analysis

Reads were mapped to the Takv6 genome (Iwasaki et al., 2021) wherein all SNP positions between Tak-1 and Cam-2 were replaced with N’s, where SNPs were identified in (Montgomery et al., 2022). Reads were trimmed with Trim Galore (https://github.com/FelixKrueger/TrimGalore). Mapping and calling of methylated cytosines was done with Bismark v0.22.2 (Krueger and Andrews, 2011). Cytones with coverage lower than 5 reads were filtered out. Replicates were merged and mapped to genes with BEDTools v2.27.1 (Quinlan and Hall, 2010) and genes were filtered to have a coverage of at least 100 reads.

### SNP data analysis

Mapped reads from CUT&RUN, RNA-seq, and DNA methylation experiments were assigned to paternal or maternal genomes using SNPSplit v0.3.4 (Krueger and Andrews, 2016). Counts for the number of reads originating from either genome were calculated per sample using SAMtools v1.9 (Li et al., 2009) and BEDTools v2.27.1 (Quinlan and Hall, 2010). The maternal ratio was determined by dividing the number of maternal reads by total reads per gene. For CUT&RUN data, only data from genes with more than ten total reads in each replicate were retained. For RNA-seq data, only data from genes with more than fifty total reads across all replicates were retained. Additionally, data from genes that were completely maternal in female Cam-2 RNA-Seq data or were not completely paternal in male Tak-1 RNA-Seq data were excluded from further maternal ratio analyses if there were at least five reads. In essence, gene_exclude= filter(snp_quality_filter,(genotype==“Cam2” & ratio<0.95 & mat+pat>=5) | (genotype==“Tak1” & ratio>0.05 & mat+pat>=5)).

The statistical analysis of CUT&RUN and RNA-seq SNPs is based on prior analyses (de la Filia et al., 2021; McDonald, 2014). For each gene and replicate, an exact binomial test was performed against the null hypothesis of Mendelian expression (*p_m_*□=□0.5). In cases where the number of reads were not a discrete count, the number was rounded down before the exact test. The non-rounded values were used for estimating the expression ratio. The p-values were Bonferroni corrected for multiple testing. Significant (adjusted p < 0.05) cases were classed, based on the ratio estimate as maternal if *p_m_*□≥□0.95, maternally biased if 0.65□≤ *p_m_*□<□0.95, paternally biased if 0.05□<□*p_m_*□≤□0.35 and paternal if *p_m_*□≤□0.05. Cases with 0.35 < *p_m_* < 0.65, as well as all non-significant results were classed as unbiased. Then each gene was tested for heterogeneity across replicates using G-tests. As in (de la Filia et al., 2021), all genes that did not show significant heterogeneity across replicates (p > 0.05) were automatically included in the final analysis.

Significantly heterogeneous genes were kept only if all replicates were all either significant or all non-significant in the exact binomial test and their ratios fell into the same category. For the kept genes, the final expression ratios were estimated as the mean of the expression ratios estimated in the two replicates separately. The expression ratios were used to categorize the genes into their final groups (maternal: *p_m_*□≥□0.95, maternally biased: 0.65 ≤ *p_m_* <□0.95, unbiased: 0.35 < *p_m_* < 0.65, paternally biased: 0.05□< *p_m_*□≤□0.35 and paternal: *p_m_* ≤□0.05). The Bonferroni corrected p-values were used to call significance of the non-Mendelian expression.

### GO term analysis

Gene Ontologies were assigned to genes from the eggNOG database (Huerta-Cepas et al., 2016) using eggNOG-mapper (Huerta-Cepas et al., 2017). GO term enrichment analyses were done using topGO v2.42.0 by Fisher test (Alexa et al., 2006) using all genes with the corresponding histone modification as background.

## Acknowledgements

F.B. acknowledges support from the PlantS and Next Generation Sequencing facilities at the Vienna BioCenter Core Facilities (VBCF), and the BioOptics facility and Molecular Biology Services from the Institute for Molecular Pathology (IMP), members of the Vienna BioCenter (VBC), Austria. S.A.M. would like to thank X. Grau-Bové and E. Axelsson for advice on statistical analyses. This work was funded in whole, or in part, by the Austrian Science Fund (FWF) grants P32054, P33380 to F.B., FWF doctoral school DK W1238 to S.A.M. For the purpose of open access, the author has applied a CC BY public copyright licence to any Author Accepted Manuscript version arising from this submission.

## Author contributions

S.A.M. conceived and designed the experiments. S.A.M. performed the RNA-seq, CUT&RUN, Bisulfite sequencing, and imaging experiments. S.A.M. performed statistical analyses, analyzed data, created figures, and curated data. F.B. supervised the study. S.A.M. and F.B. wrote the manuscript.

## Competing interest declaration

The authors declare no competing interests.

## Data and materials availability

The CUT&RUN and RNA-seq sequencing datasets generated for the current study is available in the Gene Expression Omnibus (GEO) under series GSE229110. Publicly available datasets can be accessed under the GEO SuperSeries GSE193307 and GSE229110. Source data are provided with this paper. Original images are deposited online at FigShare. DOI: 10.6084/m9.figshare.22513258. All original code has been deposited online at FigShare. DOI: 10.6084/m9.figshare.22513993.

Any additional information required to reanalyze the data reported in this paper is available from the lead contact upon request. Further information and requests for resources and reagents should be directed to and will be fulfilled by the lead contact, Frédéric Berger.

## Notes

### Competing Interest Statement

The authors have declared no competing interest.

